# Population subspaces reflect movement intention for arm and brain-machine interface control

**DOI:** 10.1101/688259

**Authors:** H. Lalazar, J.M. Murray, L.F. Abbott, E. Vaadia

## Abstract

Motor cortex is active during covert motor acts, such as action observation and mental rehearsal, when muscles are quiescent. Such neuronal activity, which is thought to be similar to the activity underlying overt movement, is exploited by neural prosthetics to afford subjects control of an external effector. We compared neural activity in primary motor cortex of monkeys who controlled a cursor using either their arm or a brain-machine interface (BMI) to identify what features of neural activity are similar or dissimilar in these two control contexts. Neuronal population activity parcellates into orthogonal subspaces, with some representations that are unique to arm movements and others that are shared between arm and BMI control. The shared subspace is invariant to the effector used and to biomechanical details of the movement, revealing a representation that reflects movement intention. This intention representation is likely the signal extracted by BMI algorithms for cursor control, and subspace orthogonality accounts for how neurons involved in arm control can drive a BMI while the arm remains at rest. These results provide a resolution to the long-standing debate of whether motor cortex represents muscle activity or abstract movement variables, and it clarifies various puzzling aspects of neural prosthetic research.

---

Motor cortex has been studied predominantly by relating single neuron responses to movement variables. Studies of population activity, in contrast, have led to significant insights about the relationship between neural dynamics and temporal features of movement^1,2^, movement preparation^3-8^, and learning^9-11^. Many experimental paradigms have been used to dissociate endpoint kinematics (e.g. cursor position or movement direction) from intrinsic motor variables (e.g. joint torques or muscle activity)^12-18^. However, due to their residual correlations in such tasks, it has been impossible to resolve debates about whether and how both sets of variables are encoded in neuronal activity. Beyond their clinical importance for neural prosthetics, motor cortical brain-machine interfaces (BMI) provide an opportunity to address this issue because they allow these two sets of variables to be decoupled completely. By recording and analyzing the responses of the same motor cortical (M1) neurons while monkeys controlled a cursor using either their arm or a BMI^19^, we have dissociated movement kinematics from arm biomechanics and revealed separate, orthogonal representations of them in population activity.

Two monkeys were trained on a 3D target-to-target reach-and-hold task, initially using one hand to control a cursor in a virtual reality setup (Fig. 1a,b). Surprisingly, when the task was switched to BMI control of the cursor, both monkeys succeeded in performing the task within 75 seconds on their very first full BMI session. The nonlinear BMI algorithm was not trained on arm-control or action-observation data but, instead, was trained *online* each session anew, based on correlations between neural activity and an inferred, desired cursor motion^19^ (SI Fig. 1). The very short time required for successfully using BMI control probably reflects training of the BMI algorithm, not substantial de novo visuomotor learning on the part of the monkey. This suggests that BMI exploits a component of neural activity that occurs naturally in motor cortex during engagement in a motor task even when the arm remains at rest. However, because of the lack of overt movement, this component should not be related to muscle activity.

**Figure 1.**
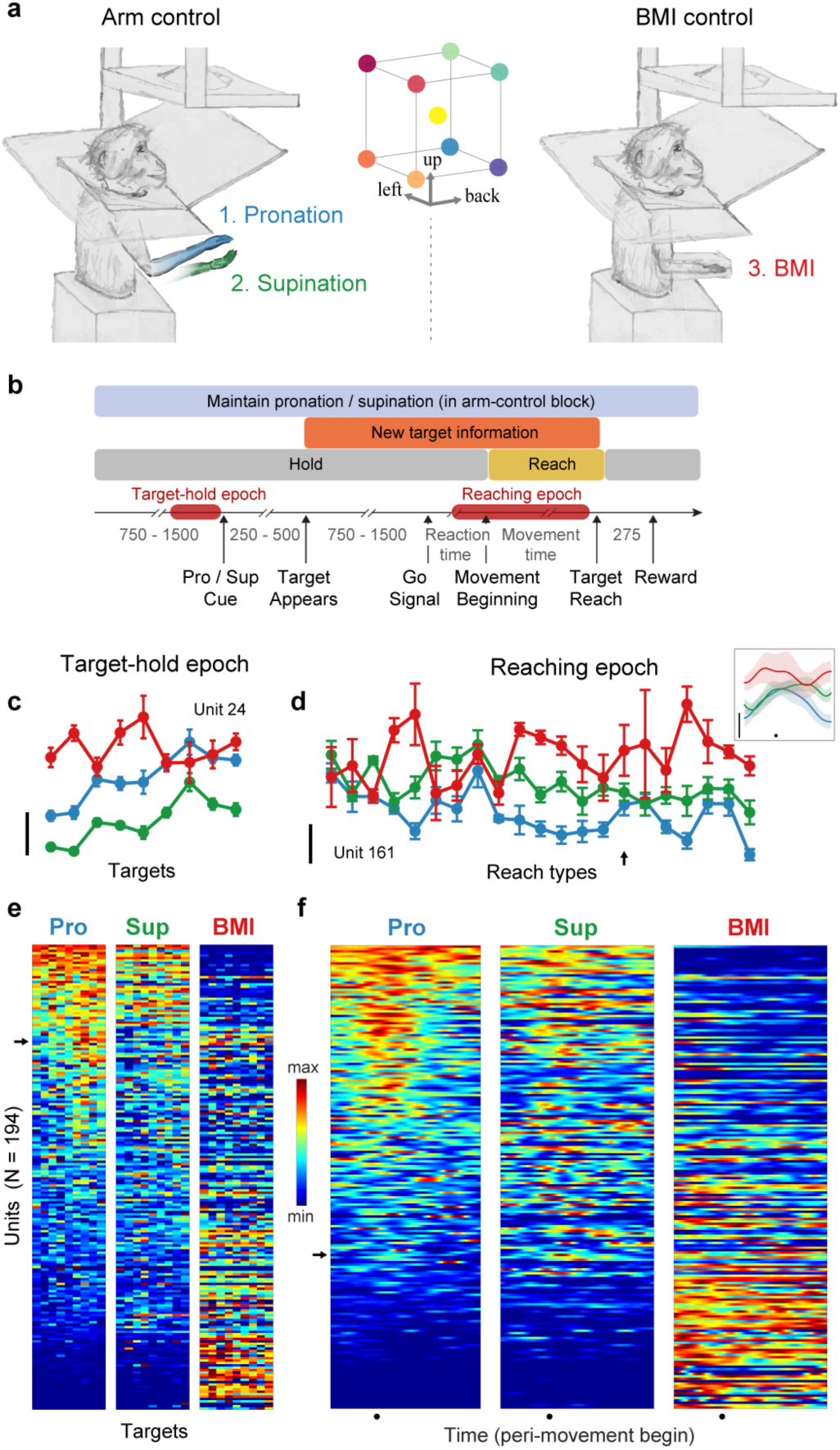
Single-neuron responses highlight the complex relationship between arm and BMI representations. **a**, Monkeys performed a continuous target-to-target reach and hold task in 3D virtual reality in one of three *control contexts:* arm control with forearm pronated, supinated, or BMI control (arm at rest). Targets appeared at 9 possible locations. **b**, Trial timeline of the task. The *Posture Cue* was an auditory tone that was different for pronation vs. supination; BMI trials had the pronation tone (which was meaningless). Double lines denote variable length intervals: randomly chosen task intervals (in ms), during *Reaction Time* and *Movement Time* due to behavioral variability. **c**, Tuning curves for example neuron during target-hold epoch show general trend: average firing rates are higher during BMI than arm control for majority of neurons, and responses are usually correlated between arm contexts (here CC = 0.97, *p* < 10^-4^) but not between arm and BMI (CC = - 0.33, -0.45, *ns*, pro vs. BMI and sup vs. BMI respectively). Mean±SEM firing rates plotted as function of nine targets held. **d**, Same as c. for another example neuron during reaching epoch. Firing rates averaged over 300 ms window around movement beginning ±SEM as function of reach types (start/target position pairs). Inset: Peri-Stimulus Time Histograms (PSTHs) for one reach type (arrow) shows deviations in firing rate and temporal profile across contexts. X-axis -200 to +300 ms around movement beginning (dot). **e**, Heat maps of tuning curves of all units during target-hold, sorted by mean firing rate during pronation only. Each row is the firing rate of one unit (arrow shows unit in c.), firing rates normalized per unit (across targets and contexts) to range 1. Normalizing obscures some significant tuning (dark blue), due to large inter-context baseline firing rate differences. **f**, Same as d. for PSTHs of all units for one reach type (arrow, d. inset). X-axis -200 to +500 ms around movement beginning (dot). Scale bars = 10 spikes/s.

We tested this hypothesis by analyzing single- and multi-units recorded during alternating blocks of arm or BMI control (hereafter called *control contexts*). In addition, the arm control context was divided into blocks in which the forearm was either pronated or supinated (Fig. 1a). Almost all recorded units modulate their firing rates in both arm and BMI control, but there is no straightforward relationship between the tuning of neuronal responses in the two contexts during either the target-hold (Fig. 1c,e) or reaching (Fig. 1d,f) epochs.

## Population activity parcellates into orthogonal subspaces representing distinct movement variables

We analyzed the structure of the neuronal population activity using a combination of dimensionality reduction and decoding techniques. We first applied principal components analysis (PCA) to the full matrix of responses across all target positions and control contexts, during the target-hold epoch (Fig. 2a; <u>SI</u> <u>Movie 1</u>). Surprisingly, the first principal component (PC), which accounts for 62% of the variance, exclusively separates arm from BMI control (Fig. 2b). The population responses projected onto this PC for the different targets in each control context overlap, so no information that could be used to direct the hand or cursor is available in this dimension. Population activity when the monkey rested his hand on the chair (during breaks in arm-control blocks) has zero projection onto (i.e. is orthogonal to) this subspace (Fig. 2b), indicating that M1 activity during BMI and resting periods is different, even though the arm is similarly at rest in both cases. Therefore, this subspace does not represent whether the arm muscles are active or not. In summary, this first PC defines a 1-dimensional *context subspace* that carries a binary signal indicating the different cortical state during arm versus BMI control.

**Figure 2.**
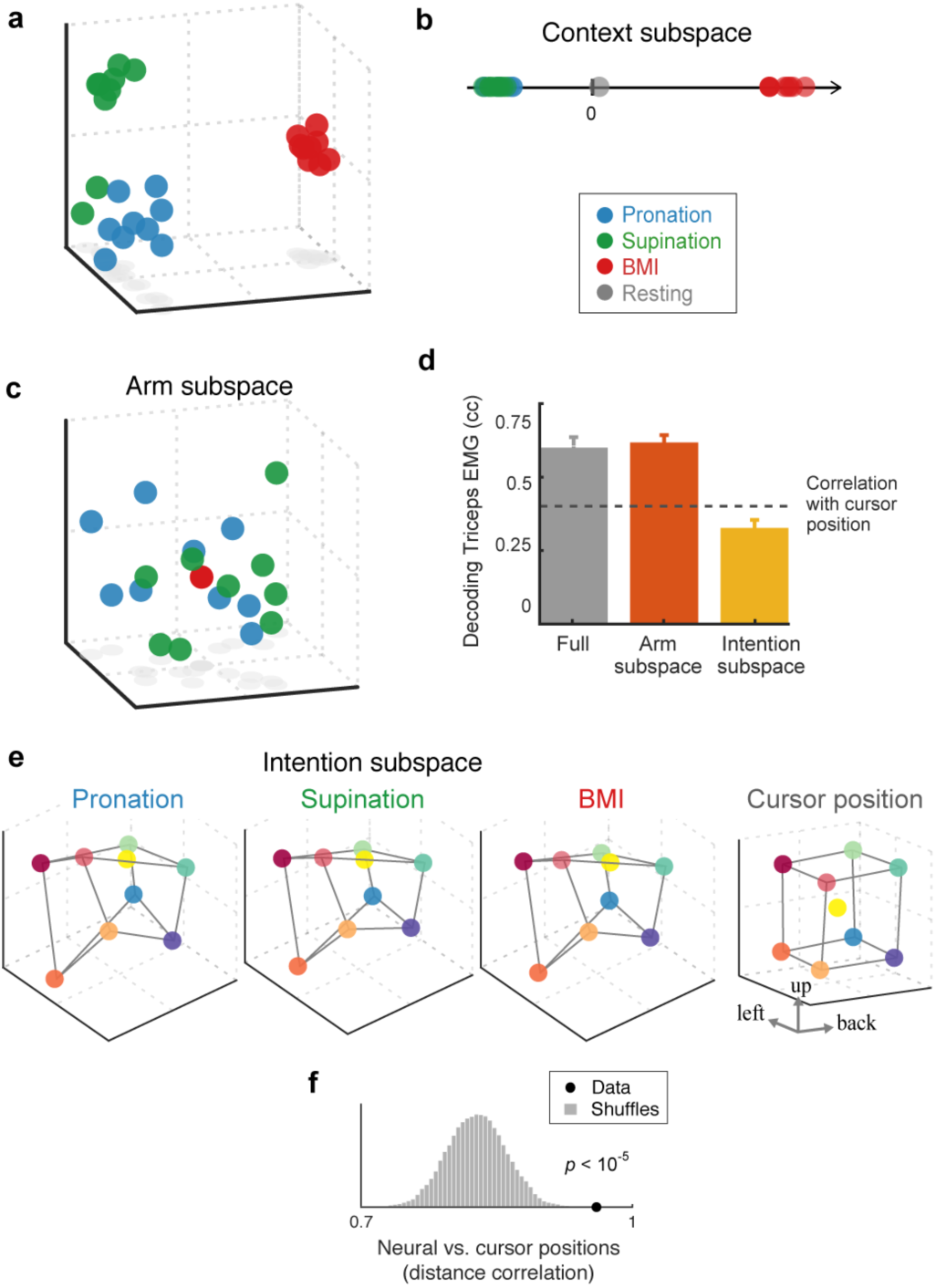
Population activity parcellates into orthogonal subspaces and reveals representation of movement intention: target-hold epoch. **a**, Population activity (Fig. 1e) projected on to first 3 principal components (PCs) (see <u>SI Movie 1</u>). **b**, Data projected on first PC shows binary separation of arm versus BMI activity, independent of targets. This *context* signal does not reflect arm at rest (gray dot), and expresses differences in baseline firing rate between contexts (SI Fig. 4). **c**, Subspace occupied only during arm control, BMI activity occupies its null-space (9 points overlap). Data projected onto first 3 (of 9) dimensions (found using a Difference-of-Covariances method; see <u>SI Movie 2</u>). **d**, Cross-validated single-trial decoding of triceps EMG shows arm subspace contains information for controlling the muscles. Median (across trials) ± 99% CI of correlation coefficient between real and predicted EMG of firing rates projected into arm subspace is not significantly different than using the full-dimensional firing rates, yet larger than the residual correlation of the cursor position and EMG, and larger than decoding using projection into intention subspace. **e**, Subspace with conditions-dependent activity that is invariant to control context, reveals an abstract “intention” representation (see <u>SI</u> <u>Movie 5</u>). Cube-like structure reflects preservation of cursor (or target) distance in external coordinates. Data projected on first 3 (of 5) dimensions (found using dPCA), accounts for 4.4% of data variance, colored by mean cursor position (right). **f.** Distance correlation between population activity in intention subspace and cursor position, for real data (dot) and shuffle controls (distribution). Neural data was averaged across contexts; shuffles were created for all (N=9! = 362,880) permutations.

The data projected onto the second and third PCs show variance during arm control but cluster around the origin during BMI (Fig. 2a). In other words, the population activity during BMI is approximately orthogonal to (or lies in the *null space* of) the subspace defined by the second and third PCs. This suggests an association of these dimensions with muscle activity. To define this association more precisely, we used a variant of PCA (a difference-of-covariances method; Methods) to find mixtures of PC dimensions that contain high variance during arm control and low variance during BMI. In the resulting 9-dimensional subspace, which accounts for 18.7% of the data variance, population activity overlaps when holding different targets during BMI control (Fig. 2c; <u>SI Movie 2</u>).

On the other hand, using single-trial firing rates projected into this subspace, we could decode the EMG of major muscles used during arm control as accurately as with the full-dimensional responses (Fig. 2d). In addition, predicted EMGs are not significantly different from zero when the same decoder is applied to firing rates recorded during BMI. Finally, training a decoder to predict EMGs during arm control and zero EMG for BMI activity does not degrade decoding performance. These analyses indicate that this subspace carries information suitable for controlling muscles, and we refer to it as the *arm subspace*.

We repeated these analyses during the reaching epoch for population firing-rate trajectories corresponding to each *reach type* (starting and target position pair) and control context. As in the target-hold epoch, a 1-dimensional subspace separates the BMI and arm control contexts (Fig. 3a,b; <u>SI Movie 3</u>), and there is an additional subspace that is activated only during arm control, with BMI activity falling in its null space (Fig. 3c; <u>SI Movie 4</u>).

**Figure 3.**
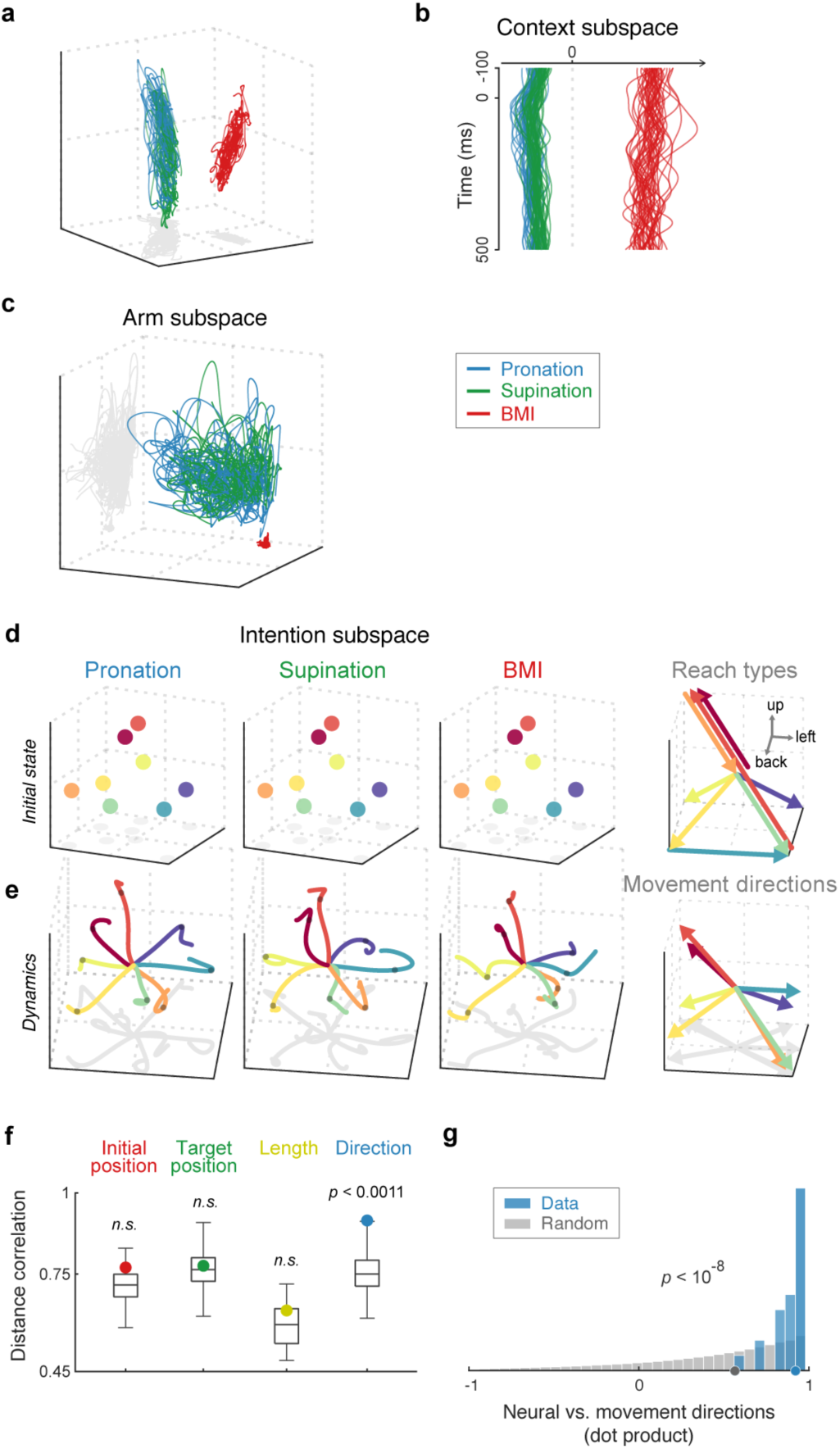
Orthogonal subspaces and intention representation: reaching epoch. **a**, Population activity (trial-averaged firing rates) projected on to first 3 PCs (<u>SI</u> <u>Movie 3</u>). One neural trajectory per reach-type (N=41, session 1), -100 to +500 ms around movement beginning. **b**, Neural trajectories, same as a., projected on context subspace (found using dPCA) as function of time, shows binary arm versus BMI signal. **c**, Arm subspace shows large neural trajectories during arm control, with no variance during BMI (first 3 of 20 dimensions shown; <u>SI Movie 4</u>). **d**, Intention subspace reveals context-invariant representation of upcoming reach type in initial state of neural dynamics (averaged over -100 to -75 ms before movement beginning). Population activity colored, here, by reach type (right; 8 reaches, data from 4 sessions; <u>SI Movie 6</u>). **e.** Neural trajectories for same reach types as d., after subtraction of initial state, show neural dynamics underlying movement intention that are reach-type-dependent but independent of effector (context). For clarity, only -100 to 100 ms around movement beginning (gray dots) plotted; first 3 (of 6) dimensions (<u>SI Movie 7</u>). **f.** Neural initial states (d.) are significantly correlated with movement directions, but not other kinematic variables, of planned reach (compared to shuffle distributions of all permutations, N = 8! = 40,320). **g.** Directions of neural dynamics are highly correlated (median dot product = 0.95) with cursor movement directions, only for 100 ms before movement beginning. Distribution of dot products between neural and movement directions is significantly different (2-sample Kolmogorov-Smirnov test) than distribution of random directions vectors (matched to data, N = 104; Methods).

## Shared subspace reveals a representation of movement intention

We next looked for neural activity that is shared between arm and BMI control and thus that could be have been used by the monkeys to move the cursor during BMI control. We used demixed-PCA^20^ (dPCA) to search for such signals by identifying mixtures of PC dimensions that account for variance across the hold targets or reach types but have minimal variance across contexts.

To our surprise, the target-hold data *demixes* almost completely (Demixing Index, *DI* = 0.99; SI Fig. 2), revealing a representation that is almost perfectly invariant across all three control contexts (Fig. 2e; <u>SI</u> <u>Movie 5</u>). We propose that this activity reflects movement *intention* because it represents each movement independently of whether the arm or BMI is used, and across the biomechanical differences between pronation and supination. In addition to these invariances, this representation has a cube-like structure similar to the arrangement of targets in the task workspace (Fig. 2f), indicating the preservation of cursor (or target) distance in neural activity space.

Neural activity up to 100 ms before movement onset is stationary and reflects the initial state of the population dynamics during reaching. This activity also perfectly demixes (DI = 1), revealing an intention subspace that represents each intended reach-type (and not hold position, as during the hold-epoch), and is again invariant to control context and arm biomechanics (Fig. 3d; <u>SI Movie 6</u>).

Applying dPCA to the entire reaching epoch also exposes an intention subspace that is context-invariant, but the structure of the neural trajectories during the reach are partially obscured by the large data variance of initial pre-movement states. Subtracting the initial states before applying dPCA results in demixing that clearly reveals an intention subspace during reaching (DI = 0.94; SI Fig. 2) in which neural trajectories provide a time-varying representation of movement intention, irrespective of control context or arm biomechanics (Fig. 3e; <u>SI Movie 7</u>). Remarkably, both the initial neural states and dynamics correlate with movement direction. The initial neural states significantly correlate with the planned movement direction, but not with starting cursor position, target position, or reach distance (Fig. 3f). Similarly, the *direction* of the neural trajectories, during the 100 ms *prior* to movement onset, significantly correlate with movement direction (Fig. 3g). Interestingly, this geometric matching of direction does not occur for the neural trajectories in the arm subspace which initially move in similar directions for all reach types, during this movement initiation period (mean dot product = 0.56; <u>SI movie 4</u>, compared to mean dot product = - 0.03 in the intention subspace; <u>SI movie 7</u>). In summary, the representations shared across contexts define an *intention subspace*, where both pre-movement and movement initiation population activity correlates with movement direction, but only until movement onset, after which the neural trajectories curve and this correlation is lost (SI Fig. 3; <u>SI Movie 8</u>).

## Controls for subspace results

Low-dimensional structure discovered in high-dimensional data, such as the neural activity considered here, can be an artifact. We verified our results in two ways: by replicating them using alternative preprocessing methods, learning algorithms, and cost functions (SI Table 1), and by statistical testing using resampling techniques, which produced *p*-values < 0.0007 and smaller (Suppl. Results).

## Recurrent neural network model suggests testable mechanisms

To gain a deeper understanding of network mechanisms that could give rise to the population representations and dynamics found in the data, we constructed recurrent neural networks (RNNs) to simulate the experimental task (Fig 4a,b). The RNN, which receives target position inputs as well as visual and proprioceptive feedback, controls 6 simulated muscles in a biophysical model arm (Fig. 4c)^21^ to perform a 2D target-to-target reach-and-hold task (Methods). The network also receives a control-context input and was trained to keep the model-arm at rest at a particular location (by the workspace side) when this input is on, even though the target inputs persist. After the network was trained to control the model-arm, a separate BMI decoder was trained to control the cursor on the basis of the RNN activity when the context input is on.

**Figure 4.**
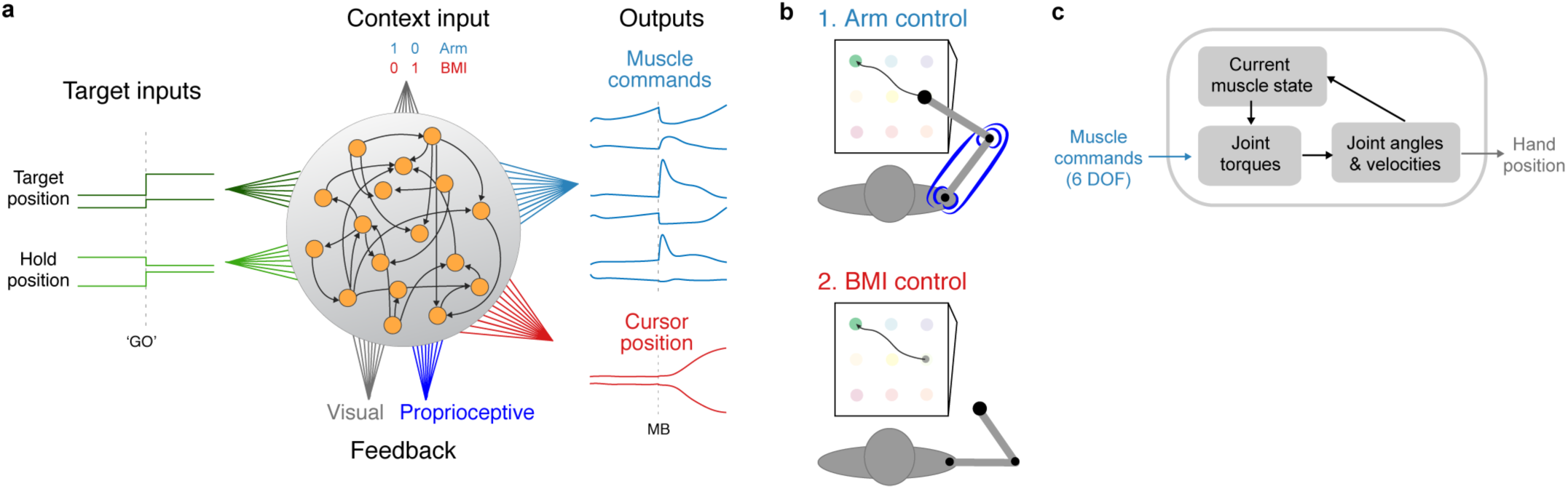
Recurrent neural network model & simulated task. **a**, Recurrent neural network model architecture. During arm-control, muscle command outputs are generated to control the model arm’s 6 muscles; during BMI control, independent readout weights control cursor directly, with arm at rest. During BMI training only BMI readout weights are adjusted. (*GO*, ‘start’ instruction, *MB*, movement beginning). **b**, Simulated reach and hold, instructed delay, point-to-point task mimics experimental task. Targets appear in virtual 2D plane, tilted at 20° angle. **c**, Muscle commands control biologically-realistic 6 DOF model arm, which includes force-length muscle dynamics, and effects of gravity.

Mirroring the rapid success with BMI control in the monkey experiment, ‘BMI’ control of the cursor by the RNN does not require any adjustment of its recurrent or input connections, but only requires training of the BMI decoder. In other words, as in the monkey experiment, the intrinsic recurrent dynamics of the RNN can be used by the BMI decoder for cursor control while the arm remains at rest. In addition to this ‘behavioral’ similarity, the activity in the RNN is remarkably similar to that seen in the real neuronal data. Analyzing the RNN’s population activity, during both the target-hold and reach epochs, exposes the same parcellation into orthogonal context, arm, and intention subspaces found in the real data (Fig. 5). Moreover, the representation in the intention subspace shows similar preservation of cursor position geometry during target-hold (Fig. 5g), and movement direction during the pre-movement (Fig. 5h) and reach initiation (Fig. 5i) periods. These results suggest that the decomposition of motor cortical population activity into these orthogonal subspaces is a ‘natural’ computational solution for motor control.

**Figure 5.**
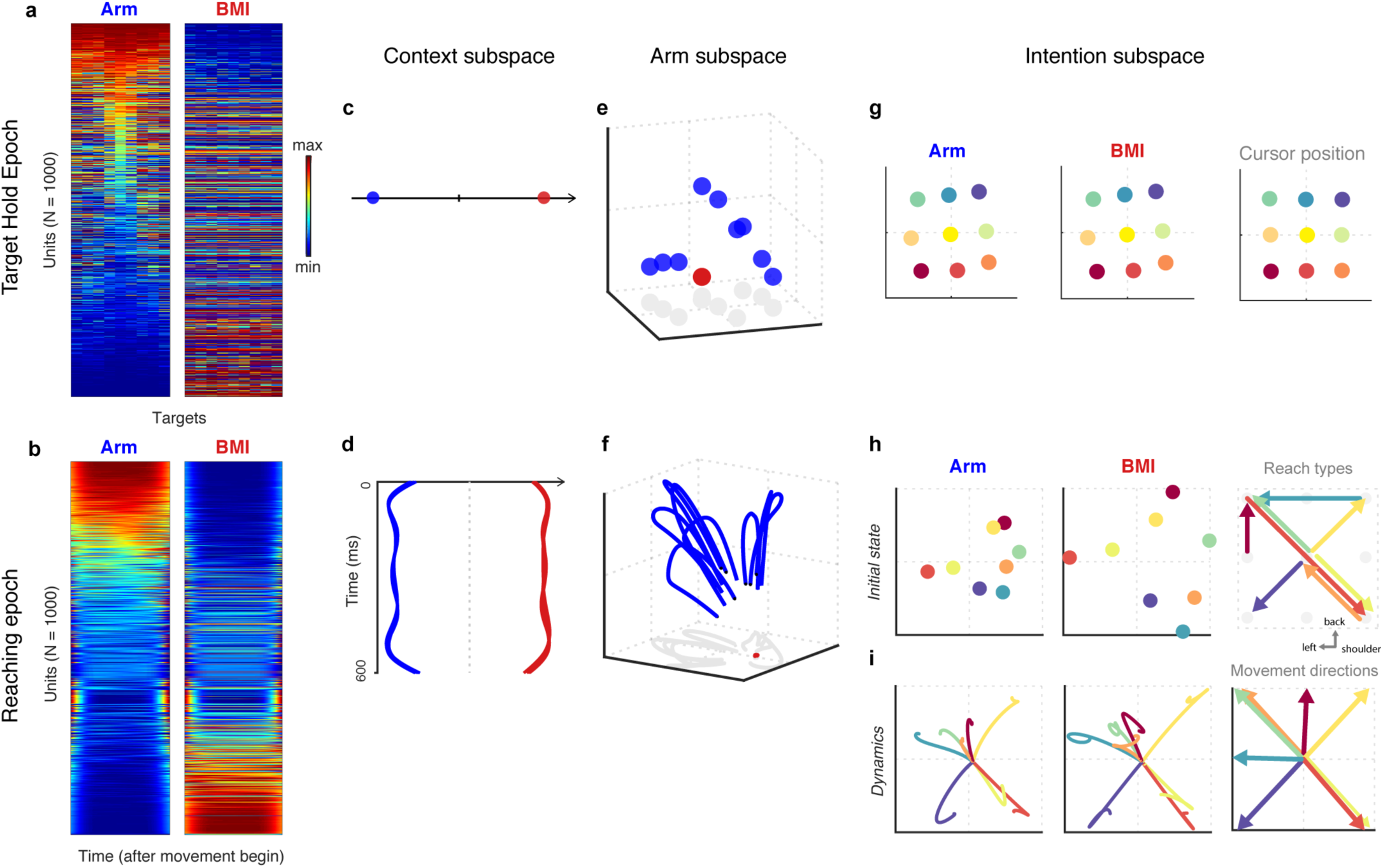
RNN activity replicates M1 population dynamics. RNN unit activity replicates the orthogonal subspaces decomposition of neuronal population activity. **a**, Compare to Fig. 1d. **b**, Compare to Fig. 1e. **c**, Fig. 2b. **d**, Fig. 3b (time after movement beginning). **e**, Fig. 2c. **f**, Fig. 3c. **g**, Fig. 2e. **h**, Fig. 3d. (reach-types are a 2D projection of reach-types in Fig. 3d). **i**, Fig. 3e.

Analysis of the RNN’s parameters enabled us to make several testable experimental predictions. In both the real and network data, the context subspace is the largest-variance component and results from changes in mean or baseline firing rates between contexts (SI Fig. 4). In the RNN, this phenomenon results from the relative strengths of the target and context inputs which, for M1 neurons, are known to arrive from distinct anatomical sources. Therefore we predict that visual target information, which M1 neurons receive via the dorsal parietal reach pathway^22^, should be approximately 7.5 times smaller than the total amplitude of the context inputs arriving to the same M1 neurons, probably from the prefrontal cortex^23,24^.

The existence of a distinct arm subspace, in both the model and data, suggests that orthogonality may serve as a selective *gating mechanism*^*6*^, allowing only certain projections of high-dimensional population activity to affect downstream structures (i.e. the spinal cord), while other projections, within the context and intention subspaces, do not. We confirmed this is the case in the network model by finding that the readout vector for each muscle (analogous to the mean synaptic weights of spinal neurons receiving motor commands) lies in the arm subspace.

## Discussion

We have shown that the nonlinear dynamics of M1 population activity segregates representations related to muscle control in one subspace from those related to an abstract time-varying *intention* of movement in an orthogonal subspace (SI Fig. 5). Each subspace relates exclusively (demixing indices close to 1) and significantly (Suppl. Results) to one set of movement variables. Furthermore, the arm subspace can be contextually ‘switched-off’ without affecting the intention subspace, as occurs during BMI control. Furthermore, this parcellation is also the ‘natural’ solution found by artificial RNNs trained to perform the task.

Our finding suggests a resolution to the enduring debate about the variables represented by M1 neurons. Single neuron responses mix signals corresponding to a number of movement variables^17,25,26^, but the population response represents endpoint kinematics and muscle-related variables in orthogonal subspaces. Mixed correlation with multiple movement variables can thus be parsimoniously explained as reflecting the relative contributions of a neuron to each of these subspaces (their coefficients contributing to each subspace’s dimensions).

Due to its almost perfect context and arm-biomechanics invariance, the intention subspace exposes an abstract population representation of movement intention. This activity is correlated with cursor (or target) position during the hold epoch^27-29^ and with movement direction during pre-movement^30^ and reaching initiation^31,32^, confirming a population representation of endpoint kinematics unencumbered by biomechanical confounds^13,31^. However, this is *not* an endpoint kinematics subspace, per se, as the cube-like representation during target-hold is consistently skewed, and the correlation during movement initiation lasts only until movement onset^17,25,32^, at which point the neural trajectories become curved and inconsistent with straight cursor movements (SI Fig. 3). One hypothesis is that M1 dynamics change at movement onset due to the integration of non-stationary feedback inputs. Our results also predict that the correlations of M1 responses during covert motor acts, such as action-observation^33,34^ and mental rehearsal^35^, reflect each neuron’s contribution to the intention subspace (Suppl. Discussion).

While the decomposition of population dynamics into orthogonal subspaces that demix distinct task variables has been found in other systems and tasks^20,24^, it remains an open question whether this is a wide-spread cortical phenomenon. This feature may relate to gating by enabling the communication of information about only selected task-related variables^6^. For example, the orthogonality of the intention and arm subspaces can account for the stationarity of the arm during BMI control even though almost all of the same neurons are active during arm and BMI control. Likewise, the orthogonality of the arm and movement-preparation subspaces may reconcile how the neurons can respond to a new target stimulus or an endpoint perturbation without affecting the arm^3,6,36^. This hypothesis could be tested experimentally by dual recordings in M1 and the spinal cord, but will have to be reconciled with the tuned preparatory activity found in spinal circuits^37^.

Confirming other studies, the *condition-independent* subspace (demixing time in other studies, or here, context) is the largest data-variance component, while the *condition-dependent* subspaces have small variance^2,20,38^. This highlights the danger of analyzing only the largest PC components, or only neurons with the largest firing rate modulations. It is unclear why networks would devote so much spiking activity to a non-coding (and in this case, binary) signal. One possibility is that, in addition to orthogonality, separation of the arm and BMI manifolds is crucial to avoid noise from causing the arm to move during covert motor acts^38,39^.

Applied to neural prosthetic research, our results suggest why decoders trained on action observation of cursor movements outperform decoders trained on data from arm movements^40-42^. Training on the arm-movement data may result in decoders that read out from both the arm and intention subspaces, but during action-observation only the intention subspace activity is likely to be present. BMIs for cursor control may benefit from identification of the intention subspace as a preprocessing step and confining the decoder to this subspace. On the other hand, limiting decoder training to the activity in the arm subspace may enhance performance of BMIs that directly control deafferented muscles^43^ or motors of robotic actuators. The immediate presence of activity in the intention subspace may explain how monkeys succeeded in using the BMI in our experiment on their very first session^19^. This provides strong motivation for developing neuroprosthetics that use online learning without pre-training (e.g. even without action-observation data)^44,45^. Across the entire reach, the neural trajectories in the intention subspace are related *nonlinearly* to cursor position (after movement onset), which may explain why nonlinear BMI decoders outperform linear ones^19,46^. This may also account for the long-term learning that occurs with daily practice when using linear BMI decoders^44,47-49^. Our results predict that this learning manifests as increased correlation between cursor position and neural trajectories in the intention subspace which, for straight cursor movements, should appear as a ‘straightening’ of curved neural trajectories.

## METHODS

### Behavioral task and Recording methods

The experimental setup, behavioral task, and recording methods were described previously^27^. Briefly, two monkeys (Macaca fascicularis) made continuous, instructed-delay, target-to-target reaching movements between 9 targets in 3D space, in a standard virtual reality setup, with their arm occluded from view (Fig. 1a). Monkeys were trained to perform the task with their forearm pronated or supinated (guided by motion capture sensors on their dorsal or palmar wrist, respectively; Phoenix Technologies), in alternating blocks of 25 trials. After behavioral training, chronic 96 microelectrodes arrays (Blackrock Microsystems) were implanted in the arm area of contralateral rostral M1. Single- and multi-unit action potentials were sorted using an automatic adaptive spike-sorting algorithm with human adjustment (Blackrock Microsystems), and offline firing-rate stability analysis was performed in a semi-supervised manner using a Hidden-Markov Model. EMG was recorded during arm-control on 6 days from monkey PK, from the Anterior Deltoid, Biceps, Triceps, Flexor Carpi Radialis, Extensor Carpi Ulnaris, using double-differential surface electrodes with pre-amplifiers x 20 (Motion Lab Systems), down-sampled, rectified, root-mean squared (20 ms window), and normalized per muscle, per day.

After several recording sessions of only arm control, each monkey completed several sessions of BMI only control. Monkey BR was first exposed to 1.5 sessions with an increasing real-time mixture of arm and BMI control of the cursor, with its arm unrestrained^19^. On the first day of BMI control for monkey PK (third, but first pure BMI control day for monkey BR), the first successful reach trial was achieved after 75 s (see BMI decoder training protocol below). Next, the monkeys performed sessions with blocks of both arm and BMI control. Data from these sessions is analyzed in this study. Visual feedback was the same for the 3 control contexts. Only single- and multi-units with stable firing rates throughout each session were included in the analysis.

Animal care was in accordance with the National Institutes of Health guidelines and was approved and supervised by the Hebrew University committee on animal experiments. The animals were housed in groups of four in an enriched playroom. Surgery involved the use of standard anesthetics, antibiotics, and analgesics that were administered pre-, during, and post-operatively as needed. Throughout the experiments, the animals were monitored daily (i.e. including rest days) and, after conclusion, were retired to a primate shelter park.

### Brain-Machine Interface

Details of our BMI decoder and training protocol were described previously^19^. Briefly, we used the Kernel-Auto Regressive Moving Average (KARMA) algorithm to predict cursor position, **y**_*t*_, in 3D: **y**_*t*_ = **W**_*ψ*_ · *ψ* (**X**_*t*:*t*–*M*_) + **W**_*χ*_ · *χ* (**y**_*t*–1:*t*–*N*_). This nonlinear decoder includes a nonlinear input-output mapping, *ψ*, operating on a history of ***M*** time bins of neural population activity, **X**, and a nonlinear autoregressive filter, *χ*, that smooths behavior, **y**, based on *N* previous time bins, and matrices of regression weights, **W**_*ψ*_ and **W**_*χ*_. Firing rate inputs were calculated with 50 ms spike counts (smoothed by a causal filter) of all single- and multi-unit channels, yielding predictions at a rate of 20 Hz. KARMA can be re-written as a Support Vector Regression problem, thus during training support vectors are accumulated (vectors of previous neuronal population activity and cursor predictions) and their associated weights are learned.

The BMI decoder was trained online, without any pre-training, i.e. at the beginning of each session all model parameters were set to zero. The initial decoder was learned on the data from the first few trials (usually 2 trials, < 20 s), during which (since the model was still empty) the cursor didn’t move. The monkeys must have generated neural activity related to their intentions to move, because this data yielded useful initial models. The trajectory ‘labels’, 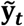, associated with the neural activity (during those trials and throughout the session) were a weighted sum of the actual cursor positions, **y**_*t*_, and the current target position, where cursor position was weighted 1 at trial beginning and 0 at its end, 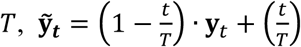. ***p***_*T*_, where ***p***_*T*_ is the 3-dimensional target position (SI Fig. 1). Decoder training was adaptive, continuing in the background throughout the session, and (after each learning iteration converged, 10-30 s) a new model was swapped in during the next inter-trial-interval. Accumulating the first full set of (3,000) support vectors took 150 s, which explains the consistent ramp up of performance at the beginning of each session and why, on most sessions, the first successful trial occurred within approximately 60 s.

### Data analysis

Our analysis focused on two task epochs. During the *target-hold* epoch (750-1,500 ms period after the reward), we analyzed single-trial firing rates averaged in a 200 ms window ending 50 ms before the first sensory cue of the next trial (Fig. 1b). Mean firing rates were averaged across trial repetitions (mean 41.9 repetitions) for each hold target and control context. Units were concatenated across sessions (N = 194), however all results were replicated per session.

During the *reaching* epoch, we analyzed a period 100 ms before-until 500 ms after movement beginning. Reach trials with outlier cursor trajectories (in terms of cursor trajectory, speed profile shape, and direction at peak speed) per control context, were excluded from analysis. After this pruning, only reach types with a minimum of 4 trial repetitions in each context were included (mean 12.4 repetitions), and as a result 16-41 (mean 25.9) reach types were analyzed per session (out of 72 total reach types). Importantly, selection was only based on behavior and not neuronal activity. Mean firing rates were calculated by averaging single trials and smoothing with a Gaussian filter (25 ms SD). Analyses were calculated per session (Fig. 3 a-c), except for Fig. 3 d-g which, in order to increase the number of neurons, used the neurons (N = 110) concatenated across 4 sessions. These results were replicated qualitatively per session as well. As a result, for these subfigures only, only the 8 reach types that met our inclusion criterion (above) for *every* session were used.

#### Dimensionality Reduction

In contrast to analyzing the population activity in each context separately and then comparing results, our approach involved applying dimensionality reduction techniques to the entire dataset in each epoch, and then analyzing the dependence of the projected data on context, movement type, time, etc. Moreover, following Cunningham and Ghahramani^50^, we decoupled finding linear projections (dimensionality reduction per se) from significance testing of structure discovered in the projected data (Suppl. Results). During target-hold, data from each context, *g*, is a 194 units x 9 targets matrix of firing rates, **X**^*g*^ (Fig. 1d). During reaching, each context’s data matrix, **X**^*g*^, is 194 units x (600 ms time x number of reach types). Principal component analysis (PCA) was applied to the concatenated matrix [**X**^pro^ **X**^sup^ **X**^bmi^] of each epoch, where mean subtraction was performed on the complete matrix. The arm subspace dimensions are calculated by an adapted version of the difference-of-covariances method described by Machens et al.^51^. Similarly to the maximum-variance derivation of PCA, solving the following loss function: *L* = *u*^*T*^(***C***_arm_ – *α****C***_bmi_) *u* + *λ* ‖*u*^*T*^*u* - 1‖ results in an set of orthogonal projections, {*u*}, where those associated with positive eigenvalues, *λ*, maximize the variance during arm control (defined as the covariance, ***C***_arm_, of the marginalized (averaged) pro and sup activity, 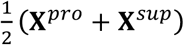 and minimizes variance during BMI, ***C***_bmi_ (*T* denotes vector transpose). The meta-parameter controlling the tradeoff between the terms, *α*, was set by cross-validation, and the results are robust to its choice (SI Fig. 6). The intention subspace was identified using demixed-PCA^20^ (dPCA), a linear dimensionality reduction method that operates on the full data-tensor which during target-hold was (*neurons*) x (*targets*) x (*contexts*), and during reaching was (*neurons*) x (*time*) x (*reach-types*) x (*contexts*). dPCA solves a reduced-rank regression problem, which seeks sets of projections of the data, each explaining the variance of only one ‘variable’ of the data (dimension of the data-tensor). For each data variable, there is no guarantee that there is even a single projection that predominantly explains its variance, so such a feature expresses an aspect of the data itself. dPCA’s regularization weight, *λ*, was selected for each application of dPCA by cross-validation, and their small values point to the negligible need for regularization with these datasets. We chose to find 26 projections for each epoch (the data rank during target hold), and verified that changing this value does not affect the original/remaining projections before the change. During the reaching epoch, we applied dPCA in three time windows, -100: 500 ms around movement beginning (SI Fig. 2), firing rates averaged (over time) in -100: -75 ms before movement beginning (Fig. 3d), and -100: 210 ms (Fig. 3e) to focus on the dynamics around movement initiation. Following^20^, we define the *Demixing Index (DI)* of a subspace spanned by the vectors {**u**}, as the mean, over the subspace dimensions, {**u**^*μ*^}, of the variance explaining only the data variable *ϕ* (dimension of data-tensor): 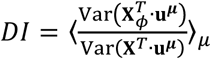, where **X**_*ϕ*_ is the marginalization of the data-tensor (defined as, the appropriately centered, average of the data across all tensor dimensions, except *ϕ*)^20^.

#### Correlations with cursor kinematics

We quantified any correlation of population activity in the intention subspace and cursor kinematics using the Distance Correlation, defined as the correlation between the distance matrices of two sets of vectors^52^. This measure allowed the comparison of neural activity and mean cursor position (Fig. 2f), initial cursor position, target position, reach length, or movement direction (Fig. 3f), without explicitly learning any parameters (e.g. as would be necessary for decoding). Median values for the shuffle controls are larger than zero, because Distance Correlation can also capture nonlinear interactions. Correlation between the *direction* of neural trajectories and movement direction was quantified as the distribution of dot-products between each pair (neural and cursor) for matching reach-types. Direction vectors were calculated using orthogonal regression (PCA), for neural directions using the data in a 100 ms window before movement beginning, and for cursor directions on the mean cursor trajectories, across trial repetitions. The set of neural directions vectors (across conditions and contexts) was rotated to best match the movement direction vectors using a single rotation matrix, learned with the Kabsch algorithm (SVD). Random control vectors (Fig. 3g, gray) were generated, for each resample, by sampling 8 random direction vectors in ℝ^3^, and then generating two more sets (for the other 2 contexts) of 8 random direction vectors, whose pairwise angles (for the same reach-type) match that of the real data. Due to the learned rotation, the median value of the random controls is larger than zero.

#### Decoding

Similar to our previous study^27^, we decoded the EMG of five major proximal muscles during the target-hold epoch. Pseudo-simultaneous population activity was created by concatenating randomly sampled single-trials from each neuron from the same *condition* (target and arm posture). Linear decoders were trained using the LASSO algorithm to decode the EMG of each muscle for 18 conditions (i.e. 9 targets in two forearm postures), and tested in a leave-one-out cross-validation scheme (1,000 repetitions). Decoding performance was evaluated, per muscle, by the correlation coefficient between the 18 predicted and real EMG values. We repeated this decoding experiment twice, once by projecting the single trials on the arm subspace, and in another experiment on the intention subspace. For population activity vector ***x***^*i*^, for single trial *i*, and subspace dimensions given by the columns of the matrix **W**, the projection calculated was **WW**^*T*^***x***^*i*^, which projects the data into the subspace but then returns it to the population activity’s original coordinates, so decoder learning will occur in the same dimensional space as in the full data decoding experiment. The subspace dimensions were learned on all the data, i.e. also on the held-out test data, which could bias the cross-validation. However, the 18 held out single-trials likely had a minimal effect on the correlation structure of the thousands of trials underlying the mean firing rate responses, and the dimensionality reduction did not use the EMG (i.e. is unsupervised). The lack of bias can be seen in the difference in decoding accuracy between the projections on the arm and intention subspaces, albeit sharing this preprocessing scheme.

### Recurrent neural network

A schematic of the recurrent neural network (RNN) model is shown in Fig. 4a. The equation describing the RNN’s dynamics is:

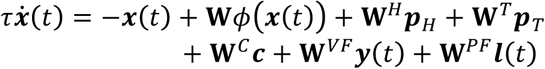

where ***x***(*t*) is an *N*-dimensional vector of RNN unit activity levels (*N* = 1000) at time *t, ϕ* is the tanh activation function, **W** is the matrix of the recurrent weights, ***p***_*H*_ and ***p***_*T*_ are the current hold and target positions, respectively (in Cartesian coordinates), **W**^*H*^ and **W**^*T*^ are their respective input weight matrices, ***c*** is a binary context input vector, with associated input weight matrix **W**^*C*^, ***y***(*t*) is the vector of visual feedback of the current (or in another version of the RNN, delayed) cursor position, with associated input weights matrix **W**^*VF*^, ***l***(*t*) is the proprioceptive feedback of current (or delayed) muscle fiber lengths and first derivatives, with associated weights **W**^*PF*^, and *τ* = 20 ms is the network time constant.

The network’s muscle-command outputs are generated by:

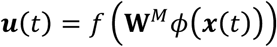

where ***u***(*t*) are the 6-dimensional muscle commands at time *t*, **W**^□^ is a readout weight matrix, and ***f***(***z***) = 0.0083 · *In* (1 + *e*^2*z*^) is a soft-rectification function. During BMI, cursor position, ***y***(*t*), is controlled directly by the following network outputs:

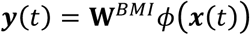

where **W**^*BMI*^ is the BMI readout weight matrix.

All input weight matrices **W**^*H*^, **W**^*T*^, **W**^*C*^, **W**^*VF*^, **W**^*PF*^ are random, sampled from the uniform distribution over [-3, 3], and fixed. The recurrent weights **W** are initialized randomly from 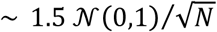 and are adjusted during training. During training, the recurrent, **W**, and muscle readout weights, **W**^*M*^, are adjusted using the Full-FORCE algorithm^53^. During training for arm-control, the context input is set to ***c*** = (1,0), but is interleaved with batches of trials with ***c*** = (0,1) in which the arm is trained to remain at rest at the side of the workspace (cursor doesn’t move). During these arm-at-rest training trials, an additional randomly fluctuating input is injected into the network to avoid it from learning a fixed point.

The model arm that is driven by the RNN is adapted from^21,54^ (Fig. 4c), and follows these forward dynamics:

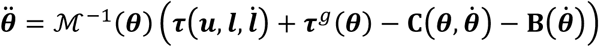

where ***θ***(*t*) is a vector consisting of the shoulder and elbow joint angles, ℳ (***θ***) is the moment of inertia tensor, and the current muscle commands ***u***(*t*), current muscles lengths ***l***(*t*) and velocities ***l***(*t*) determine the arm’s two joint torques ***τ***(*t*). The matrices **B, C**, and the dependence of the muscle torque on the other model parameters are provided in^21^. In order to require non-zero posture activity, we introduced torque due to gravity, given by:

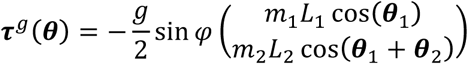

where *m*_1,2_, *L*_1,2_ are the masses and lengths of the upper arm and forearm, respectively^21^, *g* = 9,8 m/s^2^ is the acceleration due to gravity, and *φ* = 20° is the incline angle of the workspace plane.

We used the following procedures to generate labeled muscle-commands to train the network. Desired cursor kinematics were derived analytically to generate the 2-dimensional reach trajectory labels, 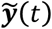, for which static cursor position during the hold-epoch is just a special case:

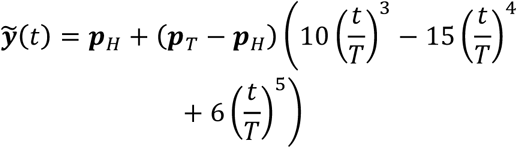

where *T* is the trial duration. This equation results from assuming trajectories are

6^th^ order polynomials, which start at the hold position, ***p***_*H*_, end at the target position, ***p***_*T*_, have vanishing initial and final velocity and acceleration, and have minimal jerk. The resulting trajectories are straight, with bell-shaped speed profiles. Inverse dynamics was solved numerically by using gradient descent to minimize this optimal-control loss function:

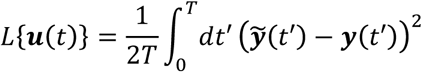

where ***y*** (*t*′) is the cursor trajectory produced by the input ***u***(*t*), assuming ***u*** 0 = 0.

During arm-control training, hold and target positions are chosen randomly within the 20×20 cm workspace. After the network has been trained, the recurrent and muscle-command output weights are frozen. Next, the BMI output weights are trained in batches, using recursive least squares. The cursor trajectory labels for the *first* training batch are the cursor trajectory labels calculated for arm-control, however for all remaining batches the labels are calculated in the same way as for the experimental BMI (as described above). Batches of training and testing continue until performance plateaus. To emphasize, during the entire BMI training phase only the BMI output weights and not the recurrent weights are trained.

## Notes

We’d like to thank M. Shachar, Y. Moshe, S. Freeman, and A. Shapochnikov for technical assistance; H. Bergman and Z. Israel for help with the surgeries. This research was supported by BSF, ISF, NIH (MH093338) grants, an NSF NeuroNex Award (DBI-1707398), and contributions from The Rosetrees Trust and The Ida Baruch Fund, and the Swartz, Simons, and Gatsby Charitable Foundations.

