## Supplementary Information for "Population subspaces reflect movement intention for arm and brain-machine interface control"

#### Supplementary Figures

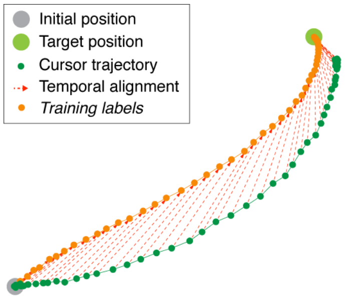


**SI Figure 1 | Trajectory labels for training BMI algorithm**. During the BMI experiment, our online BMI algorithm was trained using real-time neural activity paired with trajectory 'labels' (orange) which were a weighted mixture of the cursor position (dark green) at trial beginning and target position (light green) at trial end. See equation in Methods. This labeling technique was especially crucial for the first few trials of each session, where, because each session started with an empty model, the cursor didn't move.

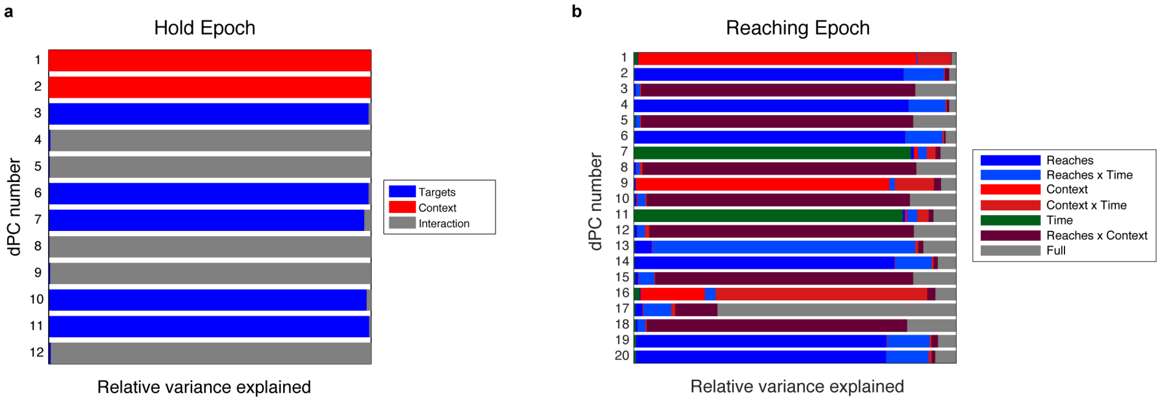


**SI Figure 2 | Relative variance explained showed almost perfect demixing in intention subspace**. **a**, Ratio of variance explained in each demixed-PC (dPC) for first 12 components during Hold epoch. Intention subspace is defined by components that demix Targets (blue), which account for this task-variable exclusively (DI = 0.99). The first component is the context subspace, replicating this result as found using PCA. **b**, Same, for the Reach epoch after subtracting the initial states (Methods). The intention subspace is spanned by components (blue) that almost perfectly demix reach-type (DI = 0.94). Again, the first component replicates the context subspace found with PCA.

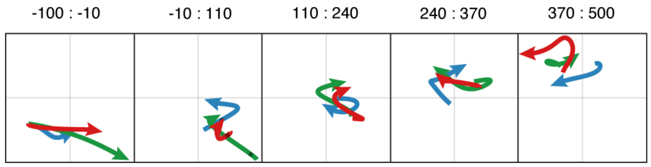


**SI Figure 3 | Intention dynamics of a single reach**. Neural trajectories for a single reach type (orange in Fig. 3d,e) in intention subspace, as a function of time (ms) peri-movement beginning (black dot). Neural activity shows divergence from correlation with cursor direction after movement beginning (see all reach types: SI Movie 8). First 2 maximal variance projections for this reach, in intention subspace, are shown.

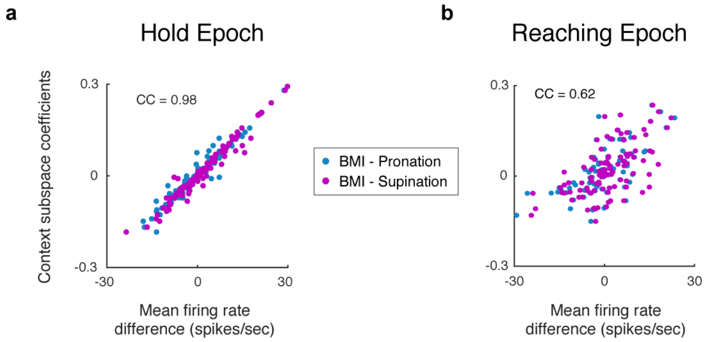


**SI Figure 4 | Context subspace results from changes in baseline firing-rates per context**. **a.** Each neuron's contribution (coefficient) to context subspace is plotted as a function of the change in mean-firing rate per context (BMI minus pronation in blue, BMI minus supination in magenta), during the target-hold epoch. Mean correlation (across both arm contexts) is shown. **b.** Same as a. for the reaching epoch. Data concatenated from 4 sessions is shown (Methods).

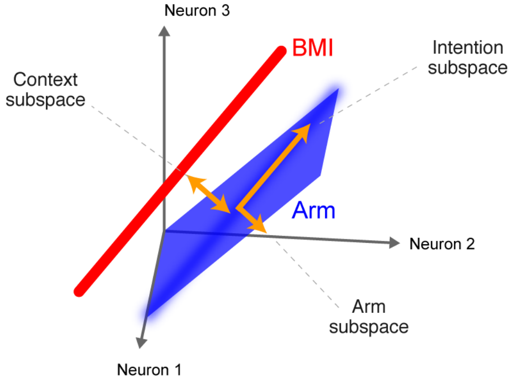


**SI Figure 5 | Diagram summarizing the relationship of arm and BMI data manifolds, across the 3 subspaces.**

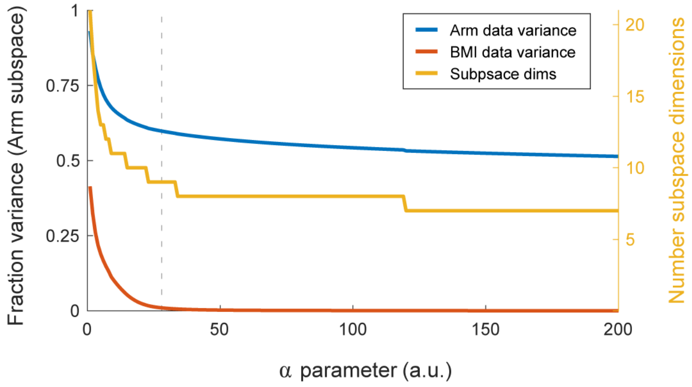


**SI Figure 6 | Robustness of difference-of-covariances α parameter.** Fraction of variance explained of the arm dataset (pronation and supination combined; blue) and BMI dataset (red) as a function of the α parameter (Methods), for one session. Curves show a large range of parameter values for which BMI data-variance is close to 0, while arm data-variance and the number of dimensions found (yellow) are stable.

### Supplementary Movies

- Movie 1: PCA - Hold epoch
- Movie 2: Arm subspace - Hold epoch
- Movie 3: PCA - Reaching epoch
- Movie 4: Arm subspace - Reaching epoch
- Movie 5: Intention subspace - Hold epoch
- Movie 6: Intention subspace, initial state - Reaching epoch
- Movie 7: Intention subspace, dynamics - Reaching epoch
- Movie 8: Arm & Intention subspace, timeline - Reaching epoch

### Supplementary Results

#### Controls for subspace results

We verified our results by replicating them using alternative preprocessing methods, learning algorithms, and cost functions (SI Table 1, below), as well as, by significance testing using resampling techniques (SI Table 2).

| | Method | Details |
|---|---|---|
| **Preprocessing** | PCA | Limit to first 12 PCs (Hold epoch), 26 (Reaching) |
| | Projection pursuit | Learn each subspace on orthogonal complement of others |
| **Learning algorithms** | PCA | |
| | Probabilistic dPCA | Brendel, et al., 2010[1] |
| | Reduced-rank dPCA | Kobak, et al., 2016[2] |
| | Unconstrained nonlinear optimization | |
| | Stiefel manifold optimization | Cunningham & Ghahramani, 2015[3] |

**Table 1:** The orthogonal subspaces were replicated using these preprocessing steps and alternative learning algorithms.

Due to the many degrees of freedom of high-dimensional data, any low-dimensional structure discovered in such data may be an artifact. We tested the statistical significance of our results following the approach of dissociating the dimensionality reduction procedure from statistical testing of the resulting low-dimensional structure found[3]. We validated each subspace result using appropriate shuffles (permutation tests) and statistics. These resampling procedures produced *p*-values for *(i)* the existence of each subspace (e.g. chance level of variance explained of that magnitude), *(ii)* defining features of the data in each subspace (e.g. to validate the Context subspace, the difference of projected data points' means, between contexts), and *(iii)* any correlation to behavior (e.g. correlation of initial state in the intention subspace and the mean movement directions), as applicable.

| | Result | Description | $p$-value | Plot | Statistic | Num resamples | Shuffle dimensions |
|---|---|---|---|---|---|---|---|
| **Hold epoch** | Context subspace (Fig. 2b) | Difference in context-dependent means | $p < 2\cdot10^{-5}$ | 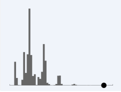 | $\langle\langle \mathbf{X}(\cdot,\cdot,g)^T \cdot \mathbf{U}\rangle_c - \langle \mathbf{X}(\cdot,\cdot,\text{bmi})^T \cdot \mathbf{U}\rangle_c\rangle_{g=\text{pro,sup}}$ | $10^6$ | Context |
| | Arm subspace (Fig. 2c) | Subspace variance explained | $p < 0.0007$ | 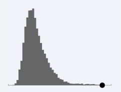 | $\frac{\text{Var}(\mathbf{X}^T \cdot \mathbf{U})}{\text{Var}(\mathbf{X})}$ | $10^6$ | Context |
| | | Arm / BMI data variance | $p < 5\cdot10^{-5}$ | 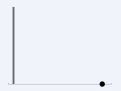 | $\frac{\langle \text{Var}(\mathbf{X}(\cdot,\cdot,g)^T) \cdot \mathbf{U}\rangle_{g=\text{pro,sup}}}{\text{Var}(\mathbf{X}(\cdot,\cdot,\text{bmi})^T \cdot \mathbf{U})}$ | $10^6$ | Context |
| | Intention subspace (Fig. 2e) | Subspace variance explained | $p < 10^{-6}$ | 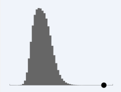 | $\frac{\text{Var}(\mathbf{X}^T \cdot \mathbf{U})}{\text{Var}(\mathbf{X})}$ | $10^6$ | Conditions |
| | | Neural states vs. cursor positions distance-correlation | $p < 10^{-6}$ | 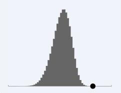 | $\text{DistCorr}(\langle \mathbf{X}(\cdot,\cdot,\text{g})^T \cdot \mathbf{U}\rangle_g, \mathbf{P})$ | $10^6$ | Conditions |
| **Reaching epoch** | Context subspace (Fig. 3b) | Subspace variance explained | $p < 10^{-4}$ | 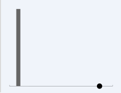 | $\frac{\text{Var}(\mathbf{X}^T \cdot \mathbf{U})}{\text{Var}(\mathbf{X})}$ | $10^4$ | Context |
| | | Maximal demixing context | $p < 10^{-4}$ | 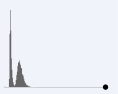 | $\max_\mu \left(\frac{\text{Var}(\mathbf{X}^T_{\text{context}} \cdot \mathbf{v}^\mu)}{\text{Var}(\mathbf{X}^T \cdot \mathbf{v}^\mu)}\right)$ | $10^4$ | Context |
| | Arm subspace (Fig. 3c) | Arm / BMI data variance | $p < 10^{-4}$ | 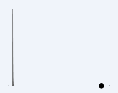 | $\frac{\langle \text{Var}(\mathbf{X}(\cdot,\cdot,\cdot,g)^T) \cdot \mathbf{U}\rangle_{g=\text{pro,sup}}}{\text{Var}(\mathbf{X}(\cdot,\cdot,\cdot,\text{bmi})^T \cdot \mathbf{U})}$ | $10^4$ | Context |
| | Intention subspace: initial state (Fig. 3d) | Neural states vs. movement directions distance-correlation | $p < 10^{-4}$ | 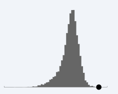 | $\text{DistCorr}(\langle \mathbf{X}(\cdot,1,\cdot,\text{g})^T \cdot \mathbf{U}\rangle_g, \mathbf{D})$ | $10^4$ | Neurons, conditions, context |
| | Intention subspace: dynamics (Fig. 3e) | Subspace variance explained | $p < 10^{-4}$ | 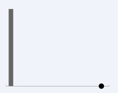 | $\frac{\text{Var}(\mathbf{X}^T \cdot \mathbf{U})}{\text{Var}(\mathbf{X})}$ | $10^4$ | Neurons, conditions, context |
| | | Maximal demixing conditions | $p < 10^{-4}$ | 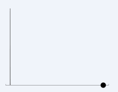 | $\max_\mu \left(\frac{\text{Var}(\mathbf{X}^T_{\text{conditions}} \cdot \mathbf{v}^\mu)}{\text{Var}(\mathbf{X}^T \cdot \mathbf{v}^\mu)}\right)$ | $10^4$ | Neurons, conditions, context |

**Table 2:** Shuffle controls for all main subspace results. Plots show distribution of statistic for shuffles (in gray) and real data's value (in black circle). $\mathbf{X}(n,c,g)$ and $\mathbf{X}(n,t,c,g)$ are mean-centered data tensors for the target-hold and reaching epochs, respectively, with indices $n$ for neurons, $t$ for time, $c$ for conditions (targets or reach-types), and $g$ for contexts (pronation, supination, or BMI). $\mathbf{X}_\phi$ is the marginalization of (data tensor's) dimension $\phi$ (see Methods); matrix multiplication involves concatenating the 2nd, 3rd (and 4th) dimensions. $\mathbf{U}$ is a matrix whose columns, $\{\mathbf{u}^\mu\}$, span the subspace stated (in each line); $\{\mathbf{v}^\mu\}$ denotes the full set of column vectors from all subspaces learned through dPCA; $T$ denotes matrix transpose. $\mathbf{P}$ is the matrix of mean cursor positions during target-hold (one row per reach-type); $\mathbf{D}$ is the matrix of mean movement directions during reaching (one row per reach-type).

#### Subspaces variance explained

The following table summarizes the percentage of data variance explained by each subspace, in each epoch. Since dPCA does not enforce the orthogonality of the subspace dimensions it finds, variance explained is calculated using the coefficient of determination method[2]. Each single context dataset (pronation, etc.) variance is calculated using mean centering from the combined data. This is done in order to include the data variance in the context subspace, which is the result of differences in context-dependent means (Fig. 2b, 3b).

| Epoch | Subspace | # dims | Combined data | Pronation data | Supination data | BMI data |
|---|---|---|---|---|---|---|
| Target Hold | **Context** | 1 | 62.3 | 37.7 | 42.9 | 86 |
| | **Arm** | 9 | 18.7 | 36 | 36 | 0 |
| | **Intention** | 5 | 4.4 | 5.6 | 5.8 | 3.1 |
| | *Other | 4, 1 | 8.4 | 11.5 | 14.1 | 3.7 |
| | Totals: | 20 | 93.8 | 90.8 | 98.8 | 92.8 |
| Reaching | **Context** | | 13 \| 37.5 | 6.7 \| 27.4 | 7.3 \| 25.1 | 23.3 \| 47.8 |
| | **Arm** | | 27.3 \| 55.9 | 33.7 \| 55.9 | 33.5 \| 66.3 | **20 \| 47.4 |
| | **Intention** | | 11.9 \| 21.3 | 12.3 \| 22.5 | 13.8 \| 19.3 | 10.6 \| 23.3 |
| | *Other | | 10.7 \| 21.6 | 12.2 \| 26.1 | 11.9 \| 22.6 | 9.6 \| 19.6 |
| | Totals: | | 51.3 \| 65.9 | 52.2 \| 64.4 | 50.8 \| 71.4 | 52.2 \| 62.6 |

**Table 3:** Percent variance explained for each subspace for the data from the Target-Hold and Reaching epochs separately. For the Reaching epoch median (left) and maximal (right) values across sessions are shown. *The cumulative variance explained for the remaining subspaces that were analyzed but, for clarity, not described in the main text. For the Hold epoch, these are (i) an additional context subspace that separates pronation from supination (but is orthogonal to BMI), and (ii) the nonlinear interaction of context and targets. During the reaching epoch, additionally, there is a subspace dependent only on time, and the full nonlinear interaction of time, reach type, and context. ** This nonzero variance is due to the within-context offset resulting from mean centering using the combined-data means (see description above; Fig. 3c).

### Supplementary Discussion

#### Intention activity, action observation, and mental rehearsal

Observation of familiar, goal-oriented, actions was found to elicit motor cortical responses resembling those that are generated when monkeys overtly performed the same action[4]. The timing of this action-observation activity *leads* the observed cursor movement and occurs even if only the task goals, but not only the cursor, were visible. This suggests the intriguing hypothesis that in contrast to “mirror neuron” activity subserving action understanding, M1 activity during action observation approximates the neural activity that generates overt action[4].

This hypothesis could account for how the monkeys succeeded in using the BMI in our experiment practically instantly: akin to action observation, they may have generated the covert component of neural activity they generate during arm control. In other words, the intention subspace could be the component of arm control activity that is generated during action observation of a familiar task. Moreover, for the first few trials in each session the cursor didn’t move (while the initial model was learned; see Methods), so only a static cursor and target were visible. For those few trials our task was identical to an action-observation experiment where only the goals are visible. The analogous question exists regarding the relation of the intention subspace and neuronal responses during mental rehearsal of motor actions (i.e. without overt action or any sensory inputs). This hypothesis can be tested experimentally by recording the same neurons while monkeys perform blocks of: *(i)* arm control, *(ii)* BMI control, *(iii)* action-observation, and *(iv)* mental rehearsal, of the same set of movements. In addition to characterizing the shared, and possibly divergent, subspaces between these four motor contexts for a well-rehearsed task, what aspects are transferred between contexts after learning a new task in only one context remains unknown. This is especially of interest given the performance improvements in overt behavior that result from motor learning by observing someone else performing the task, or by mental rehearsal.

On the other hand, M1 neural activity during BMI control (and mental rehearsal) must differ from action-observation, as the former must include an additional (minimally, scalar) input that represents *volition*—an abstract *will* for things to happen in the world. This distinguishing input is necessary for the credit assignment needed for learning—when you are only an observer and generate accurate covert activity, if the person you're observing makes a mistake you should not adapt your internal model. Additionally, neural activity during BMI must differ from both action-observation and mental rehearsal activity, since BMI experiments have shown the effects of learning to optimize control that originates from the specific small sample of cortical neurons being recorded[5,6].

#### Intention activity and eye movements

Eye movements preceding successful visually-guided reaches follow a stereotypical and well-established time course. First, the eyes complete a saccade to the target approximately at the onset of the arm's EMG activity (which precedes the beginning of hand motion by approximately 100 ms), the head then turns towards the target, and finally the eyes complete a compensatory saccade to counteract the neck movement[7,8]. For the remainder of the reach the target is foveally fixated[8,9].

Could the intention activity we described be related to eye movements? While we did not control eye movements in our experiment, their stereotypical temporal profile during reaching is inconsistent with the intention neural dynamics we discovered. Saccades to the target are initiated before the onset of the neural dynamics underlying movement initiation (~ 100 ms before movement beginning; Fig 3e), and consistent target fixation is inconsistent with the nonlinear neural dynamics in the intention subspace that occurs during the entire reach (SI Fig. 3).

1 Brendel, W., Romo, R. & Machens, C. K. in *Advances in Neural Information Processing Systems.* 2654-2662.

2 Kobak, D. *et al.* Demixed principal component analysis of neural population data. *eLife* **5**, e10989 (2016).

3 Cunningham, J. P. & Ghahramani, Z. Linear dimensionality reduction: Survey, insights, and generalizations. *Journal of Machine Learning Research* **16**, 2859-2900 (2015).

4 Tkach, D., Reimer, J. & Hatsopoulos, N. G. Congruent activity during action and action observation in motor cortex. *The Journal of neuroscience : the official journal of the Society for Neuroscience* **27**, 13241-13250 (2007).

5 Taylor, D. M., Tillery, S. I. H. & Schwartz, A. B. Direct cortical control of 3D neuroprosthetic devices. *Science* **296**, 1829-1832 (2002).

6 Jarosiewicz, B. *et al.* Functional network reorganization during learning in a brain-computer interface paradigm. *Proceedings of the National Academy of Sciences* **105**, 19486-19491, doi:10.1073/pnas.0808113105 (2008).

7 Bizzi, E., Kalil, R. E. & Tagliasco, V. Eye-Head Coordination in Monkeys: Evidence for Centrally Patterned Organization. *Science* **173**, 452-454, doi:10.1126/science.173.3995.452 (1971).

8 Biguer, B., Jeannerod, M. & Prablanc, C. The coordination of eye, head, and arm movements during reaching at a single visual target. *Experimental Brain Research* **46**, 301-304 (1982).

9 Neggers, S. F. & Bekkering, H. Gaze anchoring to a pointing target is present during the entire pointing movement and is driven by a non-visual signal. *Journal of Neurophysiology* **86**, 961-970 (2001).
